# roostR: An R package to examine diel activity patterns from Motus radio telemetry data

**DOI:** 10.64898/2026.09.10.750597

**Authors:** Kelly A. Williams, Jacob E. Morgan, Susan M. Lyons

## Abstract

1. Monitoring the diel activity patterns of free-living animals is a methodological challenge. Signal strength fluctuations from Very High Frequency (VHF) radio transmitters deployed within the Motus Wildlife Tracking System can be used as a proxy for activity. However, analytical tools to extract behavioral metrics that quantify activity patterns from these data are needed.
2. We developed `roostR`, an open-source R package that converts Motus detection data into quantitative behavioral metrics, including roost initiation and departure, roost duration, observation time, and restlessness. The package uses a sequential pipeline built around signal volatility and rolling medians to detect transitions between active and inactive states. Default parameter values were tuned using data from 55 dark-eyed juncos (*Junco hyemalis*) overwintering in southeastern Ohio.
3. We provide an example from a dark-eyed junco over a 58-day period. `roostR` estimated roost onset on 48 nights and departure on 56 mornings, with higher rolling median signal differences during the day than at night, consistent with a diurnal animal, and variable restlessness periods each night. We also used the package to estimate roost behavior of an American tree sparrow (*Spizelloides arborea*) over 43-nights.
4. `roostR` enables researchers to extract individual activity data from Motus datasets. Because all thresholds are user-adjustable, the pipeline is adaptable across species, tag specifications, and ecological contexts, enabling researchers to test hypotheses about how environmental factors influence diel activity patterns.

## 1 INTRODUCTION

The timing, duration, and frequency of activity and inactivity shape how organisms acquire resources, avoid predators, and reproduce. Diel patterns reflect selection pressures related to energy acquisition, energy expenditure, and risk avoidance (Caravaggi et al., 2018; Casula et al., 2017; Mason et al., 2017), and vary in response to photoperiod, weather, light or food availability, habitat, and predation risk (Campera et al., 2022; Caravaggi et al., 2018; Vallejo-Vargas et al., 2022). A fundamental component of diel activity is roosting behavior, characterized by the cessation of foraging and onset of sleep (Schlicht & Kempenaers, 2020; Van Hasselt et al., 2020). Variation in activity patterns among individuals and species reveals how organisms respond to environmental change (Gallo et al., 2022; Li et al., 2024; Mena et al., 2025) and informs insights for behavioral ecology, physiology, and conservation. However, continuous, non-invasive measurement of activity across free-living individuals remains a persistent methodological challenge.

Miniaturization of Very High Frequency (VHF) radio transmitters has enabled passive tracking of small animals through the Motus Wildlife Tracking System (Taylor et al., 2017) including estimation of flight trajectories (Rüppel et al., 2026). Although Motus is typically used to study movement across large spatial scales, signal strength varies with individual activity and remains stable during inactivity (Ward et al., 2013; Fig. 1), providing a passive, continuous activity proxy.

**Figure 1.**
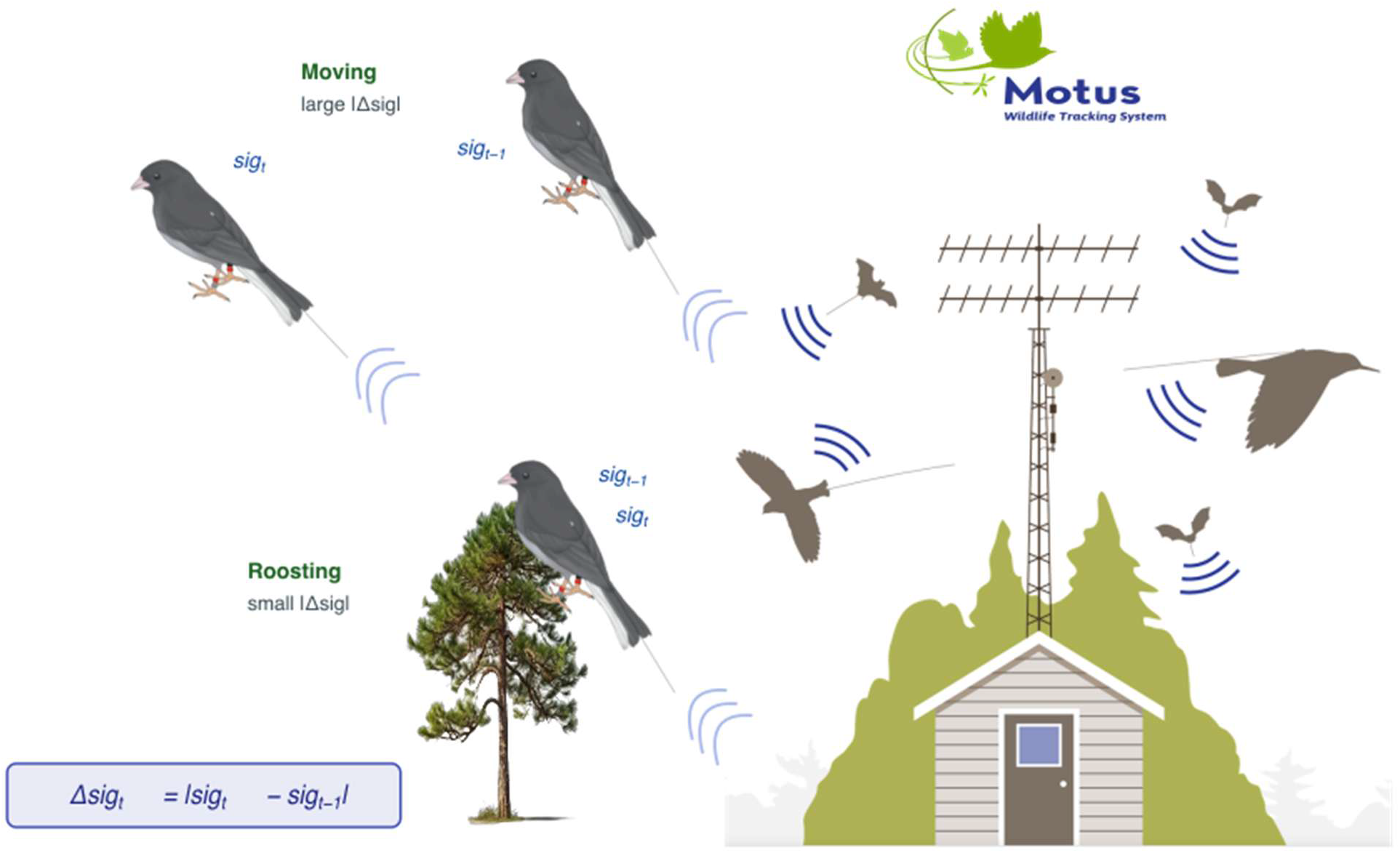
`roostR` concept. Signal strength at a Motus station varies with animal movement, increasing as an animal approaches and decreasing as it moves away from a station. An active animal will have large differences in signal strength (Δsig_t_), but a roosting/inactive animal will have small Δsig_t_. Tower and animal graphic: Motus.org, tree image: png.com, junco: JM.

Here we describe `roostR`, an R package that converts Motus detection data into behavioral metrics: onset of inactivity (roost initiation), onset of activity (roost departure), restless rate and duration of inactive periods. All detection thresholds are user-adjustable parameters, with default values tuned to data from 55 dark-eyed juncos (*Junco hyemalis*) collected using standard Motus hardware and Lotek VHF nanotags (NTQB2-5-1 and NTQB2-1; ping rates: 21.1 - 25.7 seconds).

## 2 PACKAGE OVERVIEW

`roostR` implements a sequential pipeline (Fig. 2) of five processing stages following Motus data cleaning: (1) preprocessing, (2) roost detection, (3) night observation metrics, (4) restlessness, and (5) compilation of results. Each function appends columns to the detection dataframe rather than subsetting rows, preserving the full detection record for downstream plotting and quality checks. All timestamps are handled in UTC; solar event times are converted to UTC prior to joining.

**Figure 2.**
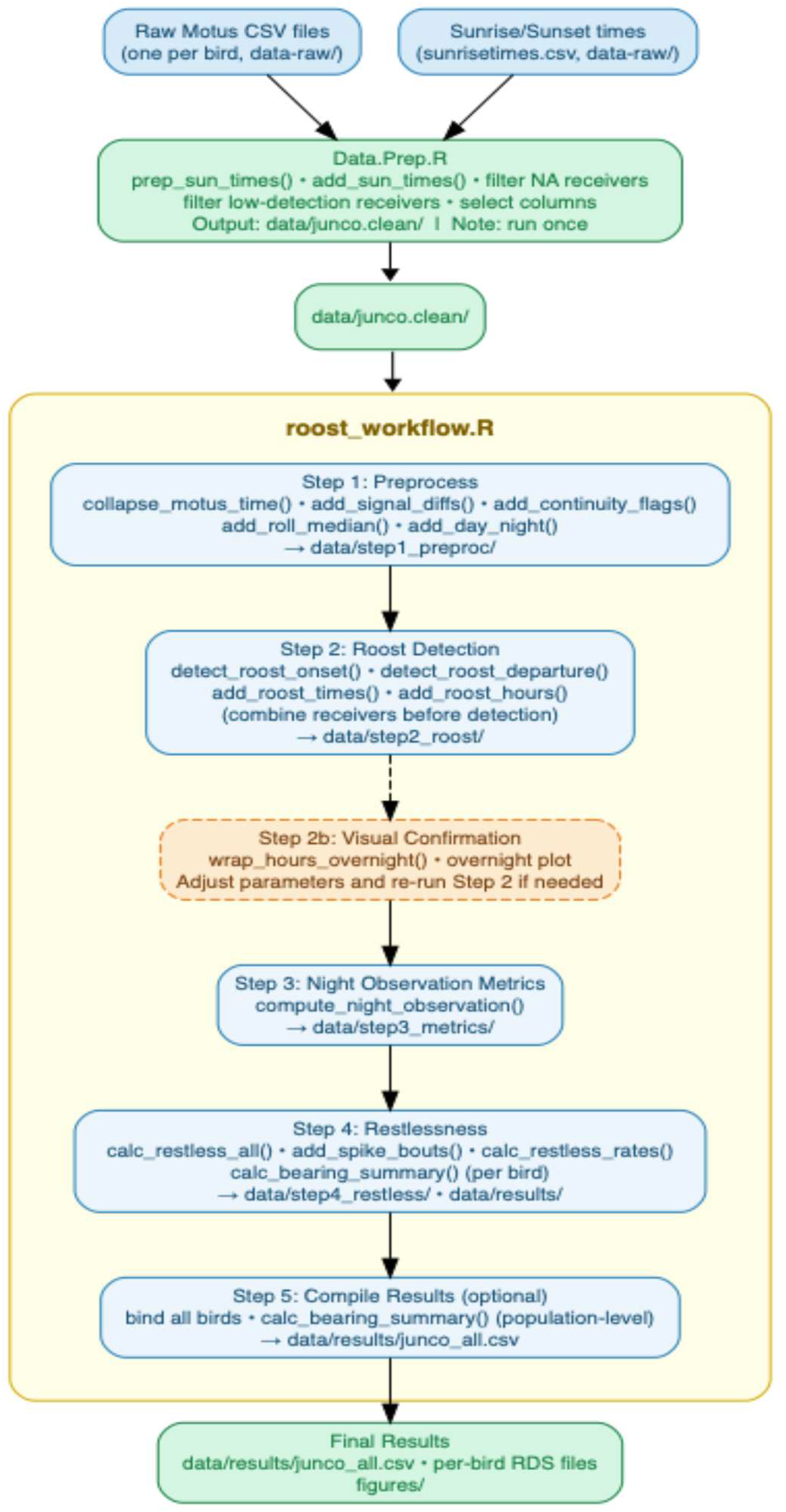
The `roostR` pipeline. Motus detections are filtered to valid detections and selected receivers. Preprocess Motus and sunrise data in Data.Prep.R before entering the main `roostR` workflow. Steps 1 - 5 are executed sequentially; output should be examined after each step, including visual confirmation of roost timing (step 2b). Final outputs include per-bird RDS files and a dataframe compiling behavioral metrics.

`roostR` requires detection data separated as one csv per individual after ambiguous or spurious detections are filtered following Birds Canada (2026) retaining columns listed in Table 1. A csv of sunrise and sunset times (or twilight values) for each location is required.

**Table 1.** Motus data required (*) and suggested for the roostR package. Year and doy (day of year) are added to the dataframe using the Data.Prep.R script.

| Variable | Type | Source | Description |
| --- | --- | --- | --- |
| tagDeployID* | Integer | Motus | Deployment-specific tag identifier; primary individual key throughout the package |
| time* | POSIXct (UTC) | Motus | Detection timestamp (UTC) |
| sig* | Numeric | Motus | Signal strength (dBm) at the receiving antenna |
| port* | Integer | Motus | Antenna port |
| recvDeployName* | Character | Motus | Receiver station name |
| runLen | Integer | Motus | Number of consecutive detections in the run |
| mfgID | Character | Motus | Manufacturer-assigned tag identifier |
| antBearing | Numeric | Motus | Antenna bearing |
| year | Integer | Data.Prep.R | Calendar year extracted from timestamp |
| doy | Integer | Data.Prep.R | Day of year extracted from timestamp |

## 3 WORKFLOW

### 3.1 Data preprocessing: collapse multi-antenna detections

A single tag pulse may be detected on multiple antennas of the same receiver or multiple receivers simultaneously if an array has overlapping detection ranges. Use `collapse_motus_time()` to group data to one row per receiver, port, and timestamp, retaining the maximum (`sig)` and the per-group mean signal strength (`sig.mean`) to ensure downstream calculations.

### 3.2 Compute activity proxies

Movement in relation to a receiving station results in variation in signal strength among time points. Signal volatility within each individual file, grouped by receiver, port combination is estimated with function `add_signal_diffs()`which returns three new variables that are core activity proxies. The absolute differences in signal strength (`sig.diff`) between consecutive detections is calculated as

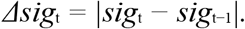

The absolute difference in mean signal between consecutive detections (`sig.diff.mean`) is calculated from `sig.mean`. The minutes elapsed since the previous detection (`time.diff`) is also calculated. Use `summary()` to examine all three variables. `sig.diff` is the primary activity proxy used in downstream behavioral detections (roost onset, departure, and restlessness), while `sig.diff.mean` feeds into a smoothing step. A stationary organism produces a near-zero signal volatility as it remains relatively fixed in relation to the receiver while movement generates larger variation in signal strength.

Gaps in the detection series are identified using a threshold based on tag ping interval using `add_continuity_flags.` Detections are marked as continuous if Δ*t* is within a user specified gap threshold (default 2-min). A new “run” identifier (`run.id`) is assigned to each uninterrupted stretch of detections and to each gap in detections. Runs provide a unit for identifying detection gaps and counting bouts for activity. Examine the proportion of detections preceded by a gap, then evaluate and adjust the threshold value.

### 3.3 Smooth signal differences

Function `add_roll_median` smooths variation in `sig.diff.mean` using a centered rolling median (default `window_min` = 15-minutes), producing a robust activity index (`roll_vol`) that is used to detect roost onset. The rolling median is computed via `slider::slide_index_dbl(),` which handles irregular detection intervals by indexing on elapsed time rather than observation count.

### 3.4 Classify detections as day or night

The function `add_day_night` creates a new variable (`diel)`which labels detection as day or night based on sunrise and sunset times (POSIXct UTC) parsed from a user-supplied lookup table by `prep_sun_times()`and joined to the detection dataframe by `add_sun_times()`. A detection at time *t* is classified as day if *t* ≥ Sunrise and *t* < Sunset; all other detections are classified as night. A diurnal organism should show increased variation in rolling median signal difference (`roll_vol`) and `sig.diff` during the day compared to at night (Fig. 3). Users should examine figures to ensure sunrise and sunset best match cessation and onset of activity and consider whether civil or nautical twilight may be better matches. Diurnal and nocturnal rolling medians should be reviewed to select parameters.

**Figure 3.**
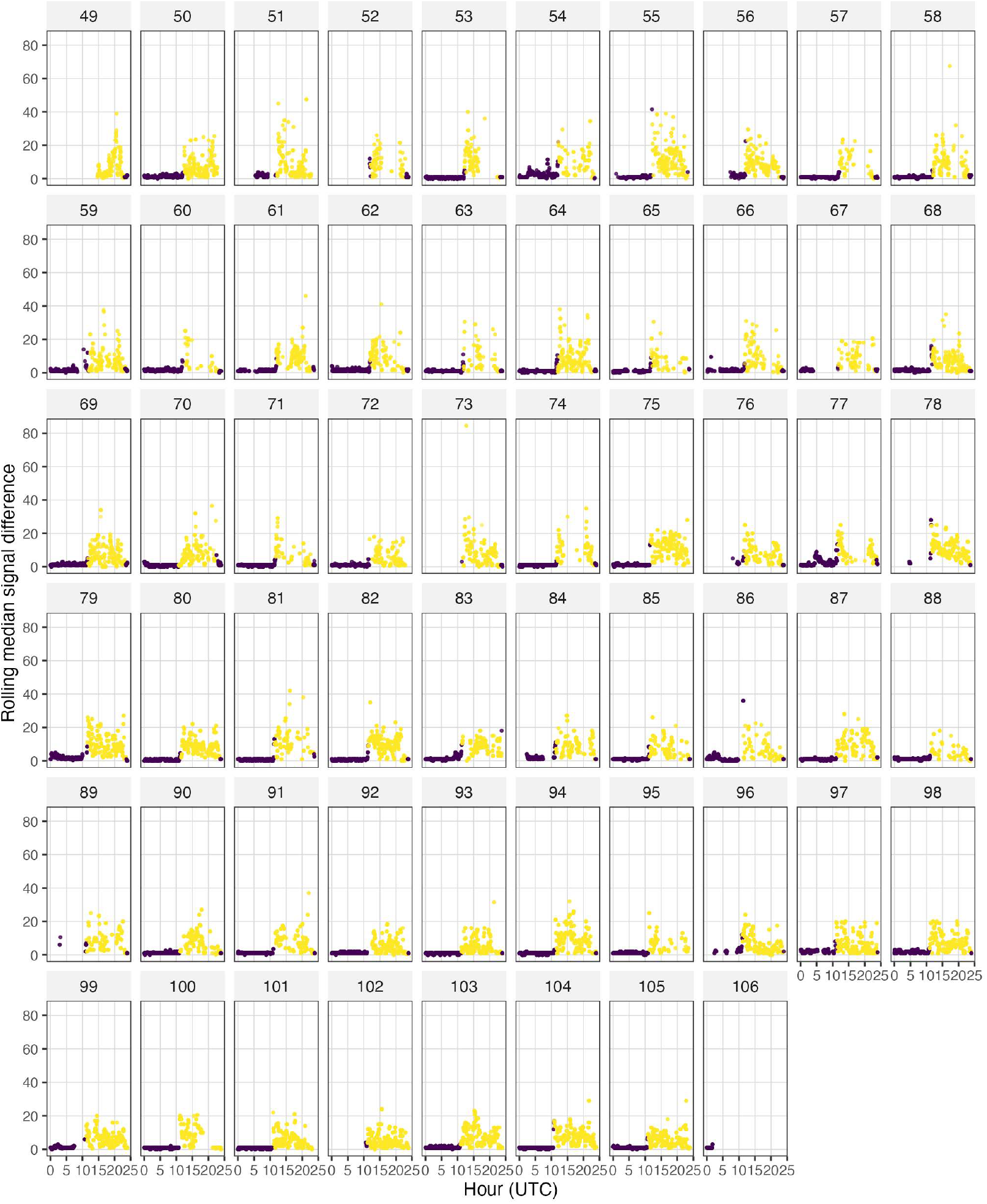
Rolling median signal difference for one bird by day of year (facet labels = Julian day) and time (UTC) with daytime values in yellow and nighttime in purple. The rolling median signal difference shows increased variation during the day when the bird moves in relation to the VHF receiving station and is relatively constant at night as the bird is relatively stationary.

### 3.5 Detect roost timing

Roost initiation is estimated using `detect_roost_onset()`. This function looks within a user defined window around sunset (default `window_minutes` = 90 min) and flags detections where `roll_vol` falls below a threshold τ (default τ = 3) as periods of low-volatility. The threshold is set between the typical nocturnal and diurnal median of `roll_vol` from the data. Consecutive low-volatility detections are grouped into runs using run-length encoding (`data.table::rleid()`). Runs shorter than a minimum duration (default `min_duration` = 10 min) are discarded. The start time of each remaining run is a candidate onset. For each candidate, the algorithm inspects the C minutes following the candidate onset (default C = 30). A candidate is accepted only if ≥ f_confirm_ (default `confirm_frac` = 0.80) of detections within this window are also low-volatility. This step guards against false positives from transient settling periods during pre-roosting activity, where variation is lower than during the day but not as low as true roosting. If detections are present in the period between the end of the confirmation window and the end of the search window, they must also satisfy ≥ f_confirm_ low-volatility detections; otherwise, the candidate is rejected. The latest candidate onset time is returned providing a conservative estimate of settlement. Candidates with no post-window detections are accepted because birds that settle low in vegetation are sometimes out of receiver range but are detected once in the air. Parameters can be set by the user to control confirmation including how long the quiet period must persist (`confirm_min`, default = 30 min) and the proportion of detections that must be low volatility (`confirm_frac`, default = 0.8).

Roost departure is estimated with `detect_roost_departure()`which identifies the time of first sustained activity each morning. The function restricts examination of detections to a window around sunrise (defaults: `window_before` = 90 minutes and `window_after` = 120 minutes to accommodate pre-dawn activity and late departures observed on some mornings). A forward rolling median of signal differences between consecutive observations (`sig.diff`) over r observations (default `roll_width` = 10) using `zoo::rollapply()` is applied to capture sustained increase in activity in relation to the receiving station instead of isolated spikes of activity associated with nocturnal restlessness. An increase in activity is flagged as roost departure if `sig.diff > δ`_s_ (default `spike_threshold` = 4) and the forward rolling median > δ_m_ (default `median_threshold` = 3) to reduce false positives. The earliest candidate departure signal is retained each morning as roost departure.

### 3.6 Visual confirmation of parameter accuracy

Roost initiation and departure in relation to sunset and sunrise and variation in signal differences should be examined visually to ensure accurate estimation of values. The function `wrap_hours_overnight()` shifts UTC hours so that 17:00 UTC (noon ET) becomes the left edge of the plot and allows the full roost interval to be visible in one panel per night. Detections are then assigned a `night_doy` with `doy` – 1 for morning detections (hour < pivot) and `doy` for evening detections so roost initiation on `doy` N and roost departure on `doy` N + 1 are treated as the same night.

### 3.7 Night observation metrics

Functions that require the complete overnight interval join departure times from `doy` N+1 back to day N via a `doy` – 1L offset applied internally. We create a biological night observation metric using `compute_night_observation()` from roost initiation to roost departure resulting in total time roosting (`time_roosting_hr`). Because VHF tags emit signals at user-defined pulse rate, and not all pulses will be detected, detection depends on Motus data filters and on the location of the animal in relation to the receiving station, resulting in gaps. Detections within this interval are assigned to time bins using `lubridate::floor_date()` (`bin_size_min` = 2 min default). Bins should be wide enough to ensure the animal is detected if present. The number of time bins with a detection is counted (`n_bins_detected`) and the amount of time an animal is observed is estimated as T_obs_ = n_bins_ × b / 60 hours. The proportion of time an animal was observed is returned as `prop_time_observed` and is estimated by p_obs_ = T_obs_ / T_roost_, where T_roost_ is the total roost-interval duration.

### 3.8 Restlessness

Restless bouts and the amount of time an organism is restless is estimated with `calc_restless_all(`). Users can define three parameters: `spike_threshold, gap_min` and `min_bout_min`. Spikes are detections where sig.diff > δ_s_ (default `spike_threshold` = 4). A new bout begins when a spike is separated from a preceding spike by more than `g` minutes (default `gap_min` = 5) or follows a non-spike detection. Bout duration is estimated as the elapsed time from the first to last spike in a bout. Single spikes are given a minimum duration to match the tag pulse interval (`min_bout_min` = 22/60). `add_spike_bouts()`annotates spikes so periods of increased activity can be observed when plotting. Then `calc_restless_rates()` uses the total time an animal was observed between roost initiation and roost departure to estimate the number of restless bouts per hour observed and the proportion of observed time during roosting spent restless allowing for comparisons among nights and individuals. Users should compare the roost duration with the amount of time observed and decide which nights to include in analyses.

### 3.9 Antenna bearing

An optional function, calc_bearing_summary(), computes the circular mean and modal antenna bearing during each roost onset interval and may be used to reveal directional roosting patterns relative to the antenna array.

### 3.10 Summary figures

Several figures should be reviewed for each individual animal using code provided in the workflow. Three example figures are provided including roost initiation in relation to sunset, roost departure in relation to sunrise and the proportion of observed time restless by day of year.

### Validation

We validated signal difference and rolling medians as thresholds using a controlled field test in which a research assistant secured a VHF tag on a board within range of a Motus station while manually recording the start and end time of each activity state (walking or stationary; n = 1,396 detections). Signal difference was lower during stationary periods (0.27 ± 0.01) than walking (22.6 ± 1.22; Fig. S1) and rolling median signal difference followed the same pattern (stationary: 0.12 ± 0.01; walking: 14.82 ± 0.62; Fig. S2). All stationary detections had a rolling median ≤ 2, supporting the use of these thresholds to distinguish active from inactive states.

## 4 IMPLEMENTATION

`roostR` was developed and tested in R (R Core Team, 2026) versions 4.5.2 and 4.6.0 and is available at https://github.com/ecologykelly/roostR (version 0.2.0, MIT license). Installation requires R ≥ 4.1.0. Core dependencies are `dplyr` (data manipulation), `lubridate` (POSIXct arithmetic), `slider` (time-indexed rolling statistics), `zoo` (observation-count rolling statistics), and `data.table` (run-length encoding). No compiled code is used; the package passed R CMD check with zero errors and zero warnings on macOS, Windows and Ubuntu.

Default thresholds were selected by evaluating data from dark-eyed juncos overwintering in southeastern Ohio, USA, including the dark-eyed junco used in the vignette sample dataset (`sparrow52550`), and refined using a larger dataset (n = 54 dark-eyed juncos, Jan-Apr 2025; xxxx et al., in prep).

### 4.1 Research Ethics

All animal handling and tagging followed North American Banding Council best practices and were conducted under United States Geological Survey bird banding permit 23434, state permit BB230012 and Institutional Animal Care and Use Protocol 00000075.

### 4.2 Example application

We demonstrate the `roostR` pipeline using two species: the dark-eyed junco dataset bundled with the package and an American tree sparrow, both detected at Motus towers in Athens County, Ohio. The worked example for the junco, with all diagnostic plots, is provided in the vignette(“roostR-workflow”). Default function parameters were used unless noted.

The package collapsed data to one detection per timestamp per receiver port combination (Table 2). Signal differences showed a clear diurnal pattern in both species (e.g., Figure 3). Default function values performed well for junco activity; however, based on daily activity patterns (Figs. S3, S4) and a lower rolling median (Table 2), threshold values for roost onset were adjusted for the tree sparrow (confirm_min = 60 min, vol_threshold = 2.5, min_duration = 15 min, confirm_frac = 0.85). The junco roosted 42 ± 4 minutes before sunset (n = 48 nights) and became active 19 ± 3 minutes before sunrise (n = 56 mornings; Figs. 4, 5). The tree sparrow became active 18.07 ± 0.08 minutes before sunrise (n = 38 mornings; Fig. SI5); however, roost onset (n = 43 nights) relative to sunset differed between standard time (2.52 ± 0.09 min) and daylight-savings time (53.61 ± 0.16 min; *F*_1, 146,224_ = 76,520, *P* < 0.001; Fig. SI6). Roost interval duration and restless activity varied among nights for both species (Table 2).

**Figure 4.**
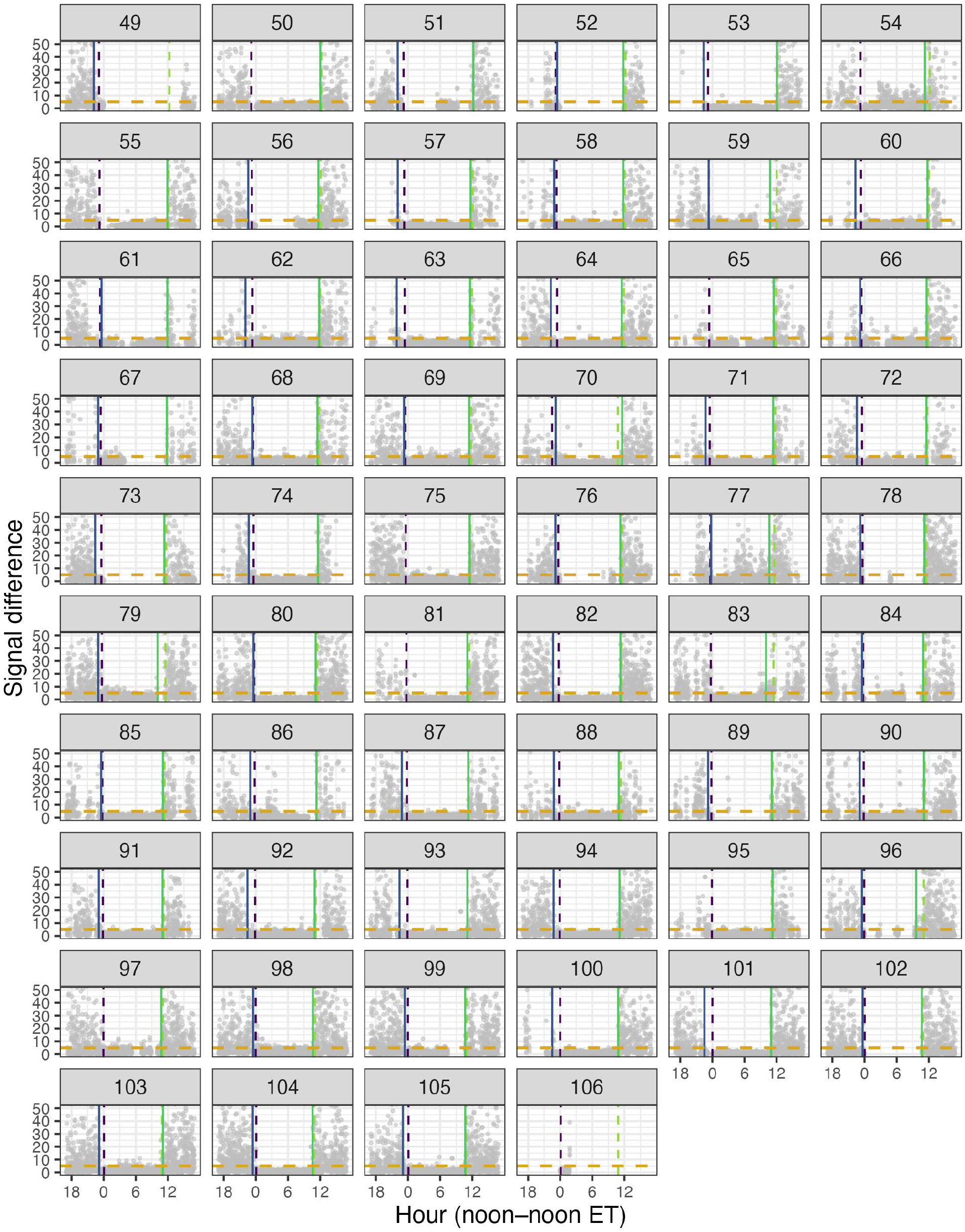
Diel activity of a dark-eyed junco by Julian day (facet labels) plotted overnight (noon to noon ET). Grey points: absolute difference in signal strength between consecutive detections (sig.diff) showing increased variation during daylight hours and low values during the roost interval. Solid lines: roost onset (purple) and departure (green). Dashed lines: sunset (purple), sunrise (green), and activity threshold for departure and restless events (yellow).

**Figure 5.**
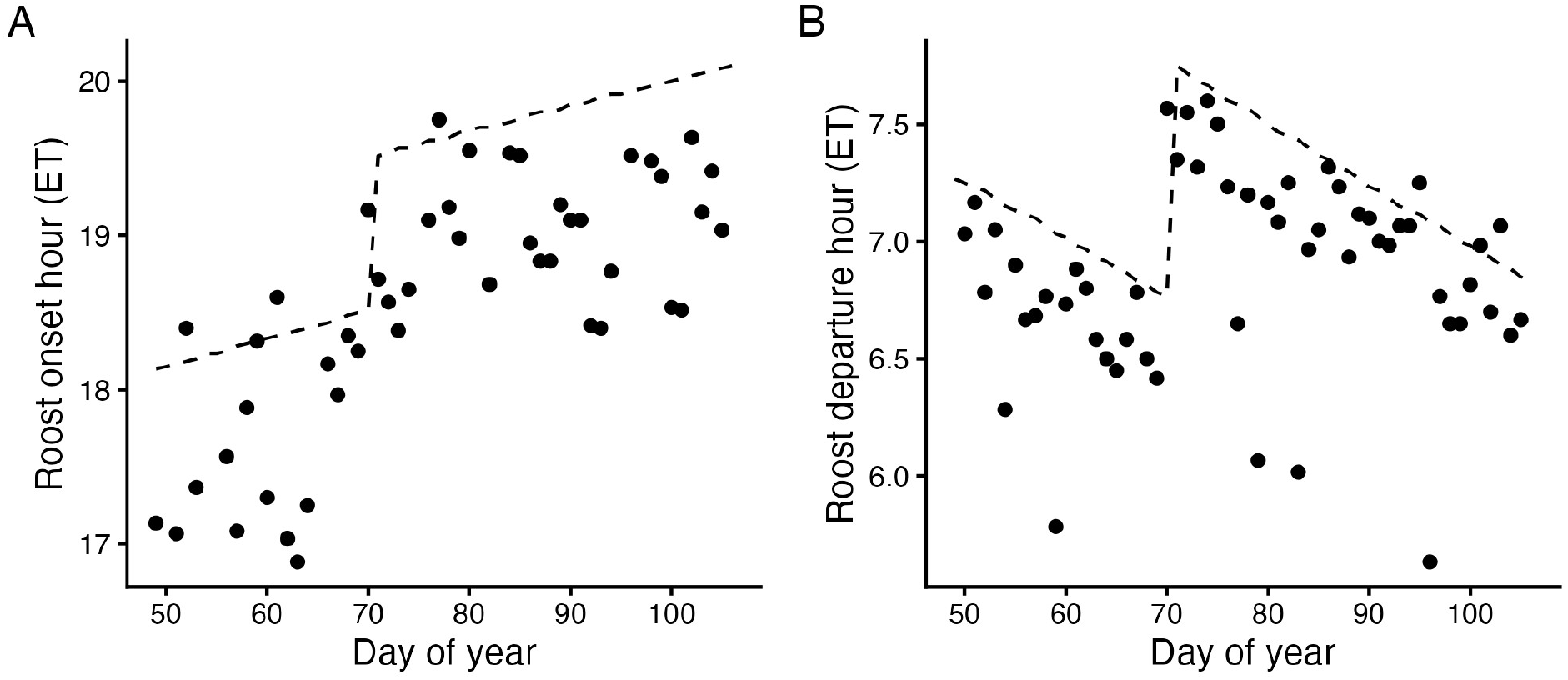
Per-night estimates of roost onset (A) and departure (B) in relation to sunset and sunrise (dashed line) by doy for dark-eyed junco.

**Table 2.** Data and package output for example application of roostR.

| Parameter | Dark-eyed Junco<br>(sparrow52550) | American Tree<br>Sparrow |
| --- | --- | --- |
| Date | 18 Feb-15 Apr 2024 | 11 Feb–26 Mar 2024 |
| Total detections | 307,978 | 168,599 |
| Non-overlapping detections | 55,233 | 168,292 |
| Rolling median | day: $7.55 \pm 0.04$,<br>night: $1.27 \pm 0.008$ | day: $1.63 \pm 0.001$,<br>night: $0.29 \pm 0.001$ |
| Roost interval duration (h) | $12.3 \pm 0.11$ | $12.75 \pm 0.11$ |
| Roost interval proportion | $0.57 \pm 0.03$ | $0.72 \pm 0.06$ |
| Average restless bouts | $15.9 \pm 2.45$ | $16.87 \pm 1.69$ |
| Total restless time (min) | $12.6 \pm 2.80$ | $6.89 \pm 0.66$ |
| Proportion time restless per observed hour | $0.036 \pm 0.007$ | $0.02 \pm 0.0001$ |
| Restless bouts per hour | $2.95 \pm 0.42$ | $3.54 \pm 0.7$ |

## 5 Conclusion

The `roostR` package provides a reproducible pipeline for extracting behavioral metrics from Motus detection data without the need to recapture tagged individuals, allowing researchers to investigate and test hypotheses about diel activity including roost timing, roost duration, and restlessness in relation to environmental contexts.

The species-specific threshold adjustments required for the American Tree Sparrow relative to the dark-eyed junco are consistent with differences in daily movement patterns (Figs. S7), and habitat specialization (Gardiner et al., 2019; Morgan & Williams, 2025). These patterns highlight that default thresholds in roostR should be evaluated against species-specific behavior, and that the diagnostic visualization workflow is essential for confirming threshold performance across taxa.

The difference in tree sparrow roost onset relative to sunset between standard time and DST warrants further investigation. Tree Sparrow movements during winter are consistent with a circadian system sensitive to twilight cues as spring approaches (Morbey et al. 2025). Sensitivity may reflect sub-Arctic breeding range, where rapid day length changes can alter circadian entrainment and activity onset (Aschoff & Wever, 1966; Reierth & Stokkan, 1998). While additional studies are needed to elucidate our findings, there is potential for roostR to compare intra- and inter-specific responses to environmental conditions.

`roostR` is an open source package that provides a framework with user-adjustable parameters to extract consistent and comparable behavioral metrics across studies, and taxa maximizing the scientific return on tagging projects within the Motus network and providing additional insights into movement and behavioral ecology.

