## Supplemental materials for "roostR: An R package to examine diel activity patterns from Motus radio telemetry data"

Supporting Information for roostR: An R package to examine diel activity patterns from Motus radio telemetry data

Validation Study

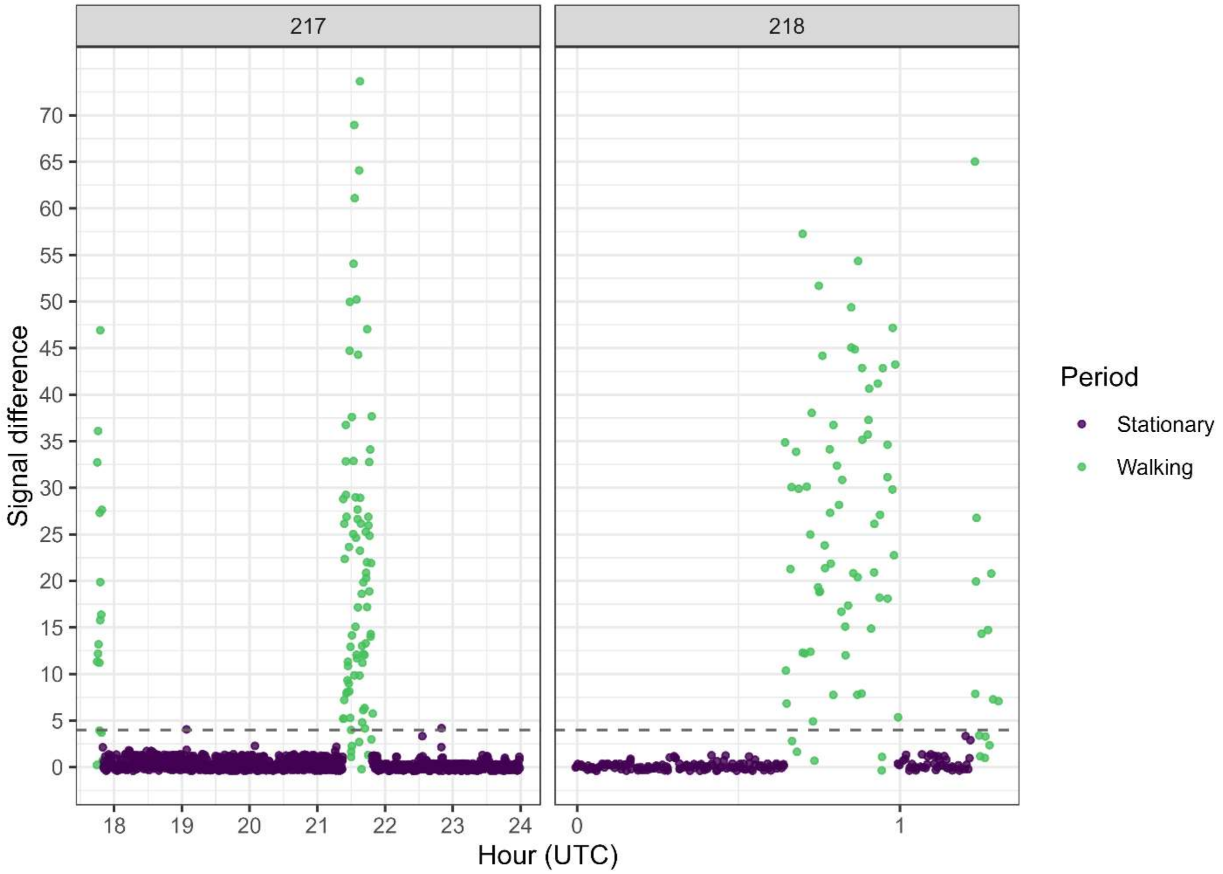


Figure S 1. Signal difference recorded during a controlled field validation. Within range of a Motus station, a VHF nanotag was secured to a board during stationary periods and carried by a research assistant during walking periods. The dashed horizontal line indicates the default signal difference threshold (≤ 4) used by roostR to classify diel activity states. Facet label = Julian day.

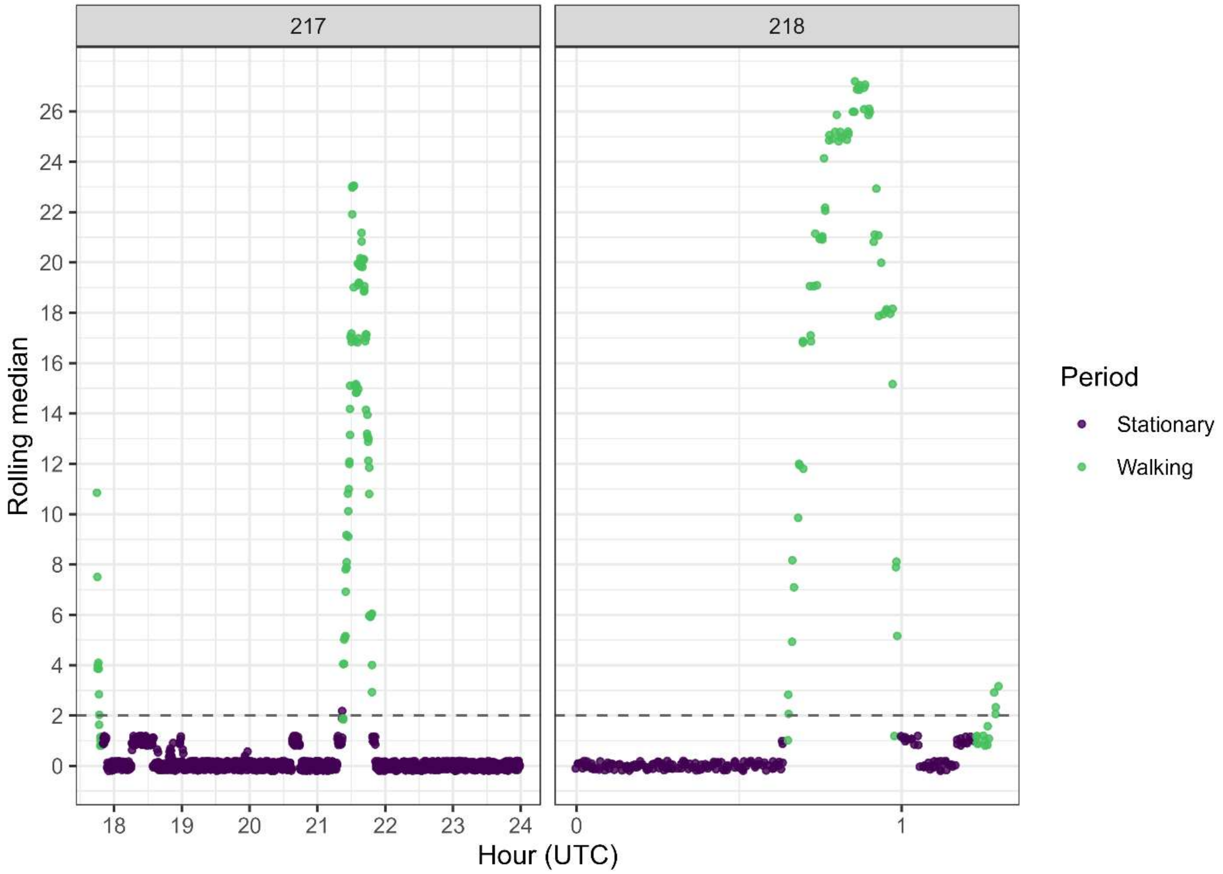


Figure S 2. Rolling median recorded during a controlled field validation. Within range of a Motus station, a VHF nanotag was secured to a board during stationary periods and carried by a research assistant during walking periods. All stationary detections fell at or below the dashed horizontal line (rolling median = 2) and 92.2% of walking detections met or exceeded this value. The rolling median signal difference is used in roostR to classify diel activity states including roost onset and departure; threshold values are adjustable within roostR functions to accommodate project specific conditions. Facet label = Julian day

American Tree Sparrow

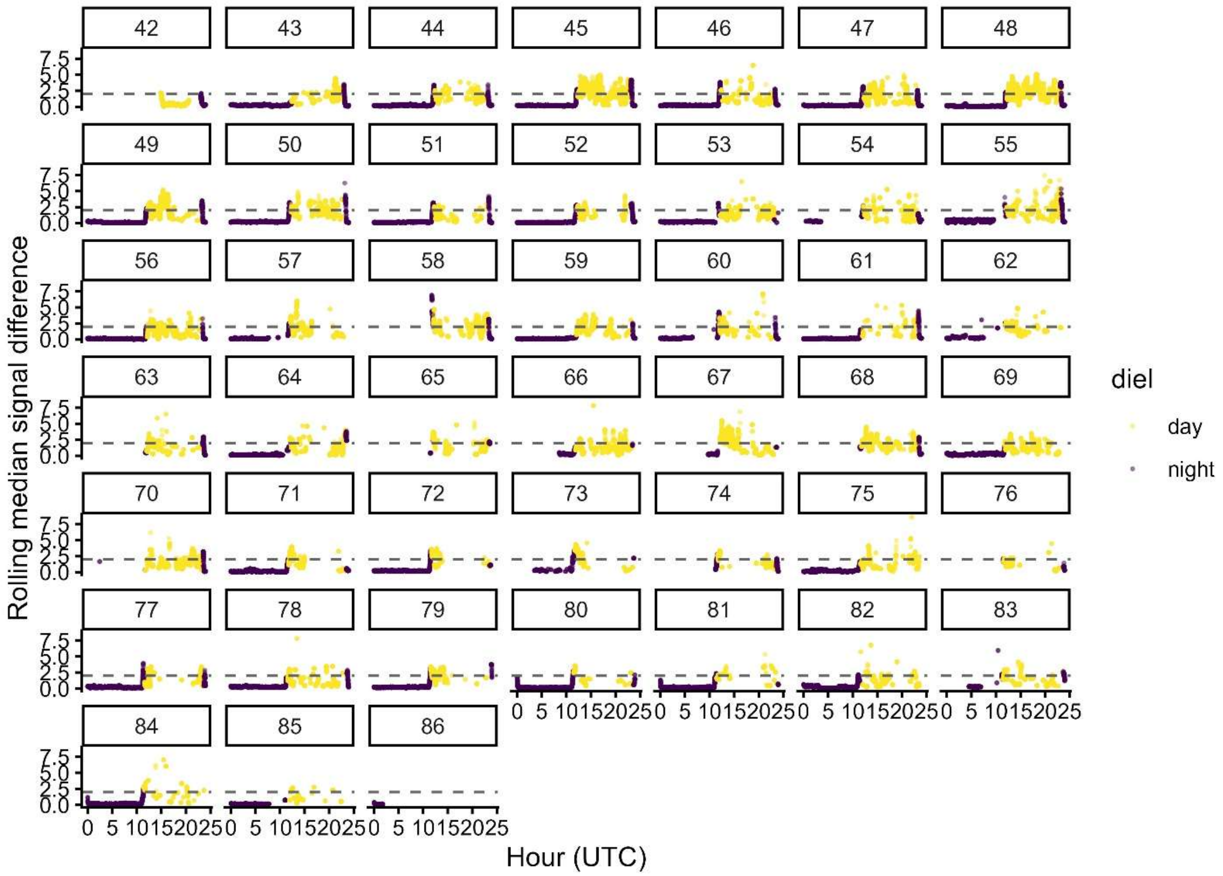


Figure S3. Rolling median during day and night for one American tree sparrow from Julian day 42 through 86 (facet labels). The horizontal line (y = 2.5) represents the rolling median threshold used in detecting roost onset.

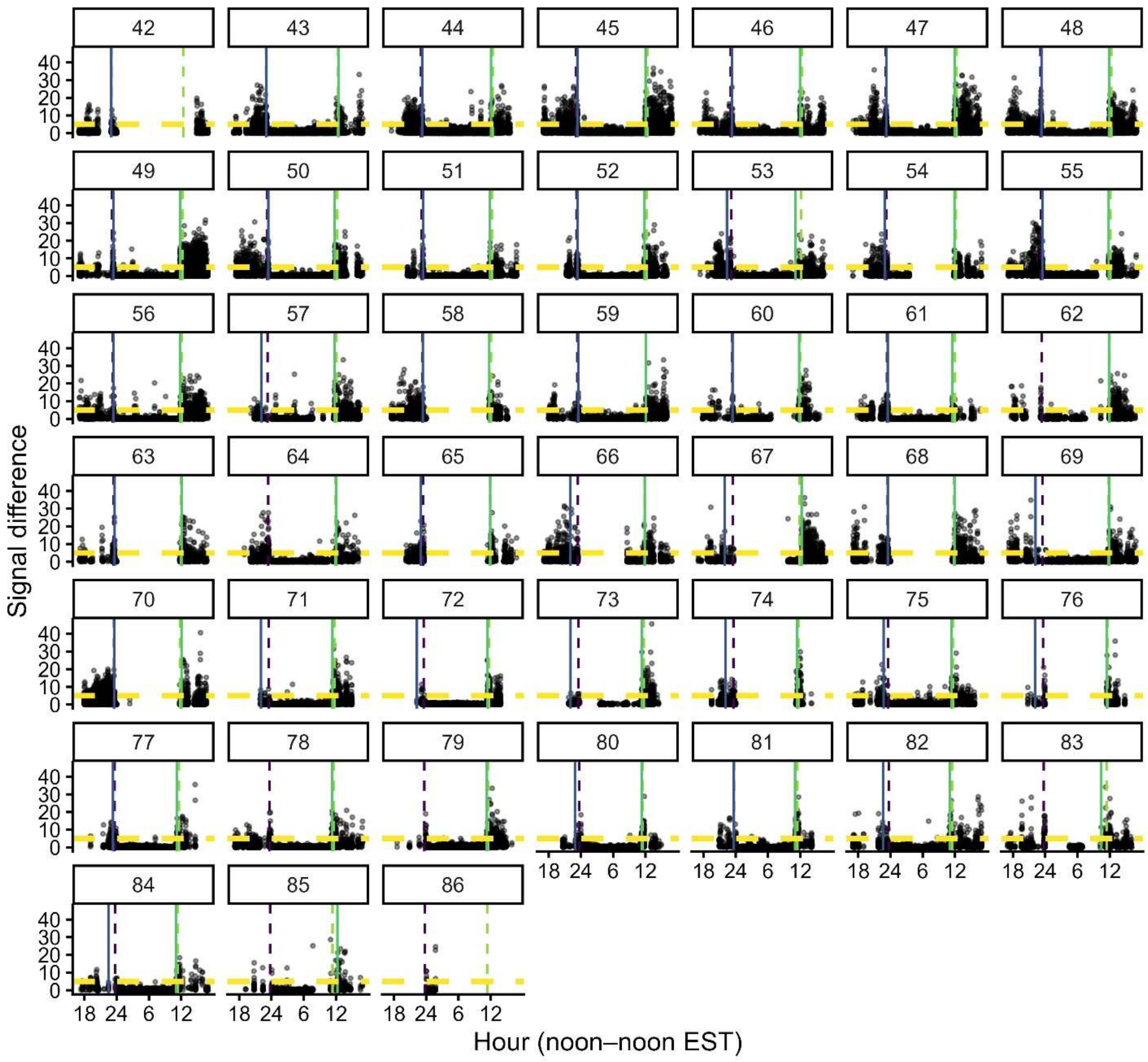


Figure S4. Diel activity patterns of an American tree sparrow by Julian day (facet labels) from noon to noon EST. Points show the absolute difference in signal strength between consecutive detections (sig.diff) for each overnight period. Solid lines indicate estimated roost onset (purple) and roost departure (green). Dashed lines indicate sunset (purple), sunrise (green), and the activity threshold for departure and restless events (yellow horizontal).

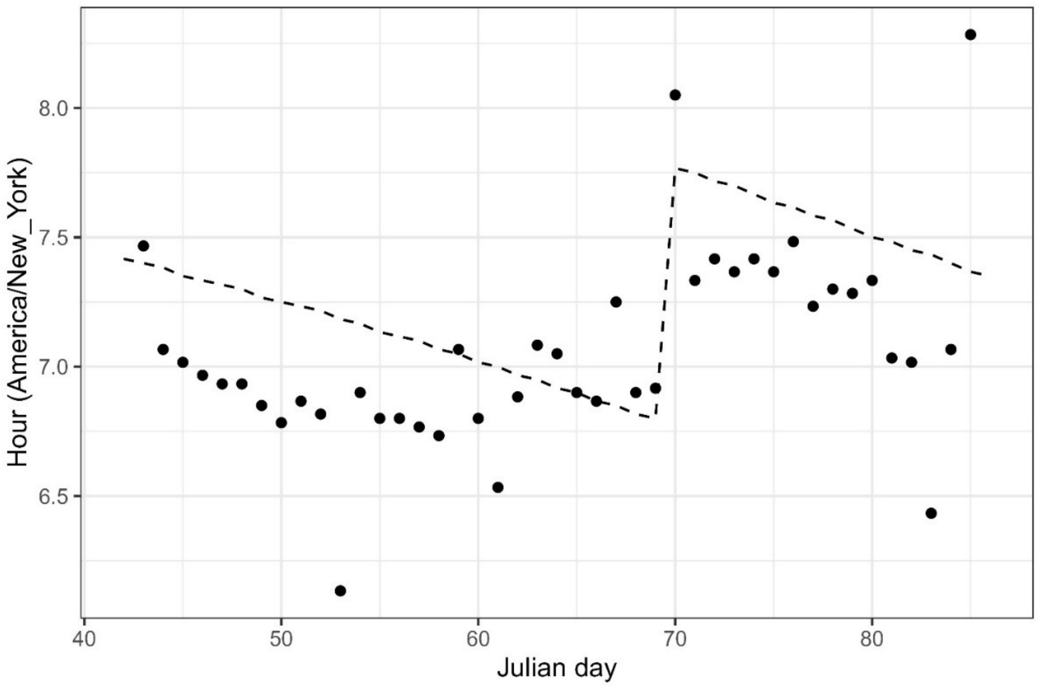


Figure S5. Per-night estimates of roost departure in relation to sunrise (dashed line) by doy for the American tree sparrow.

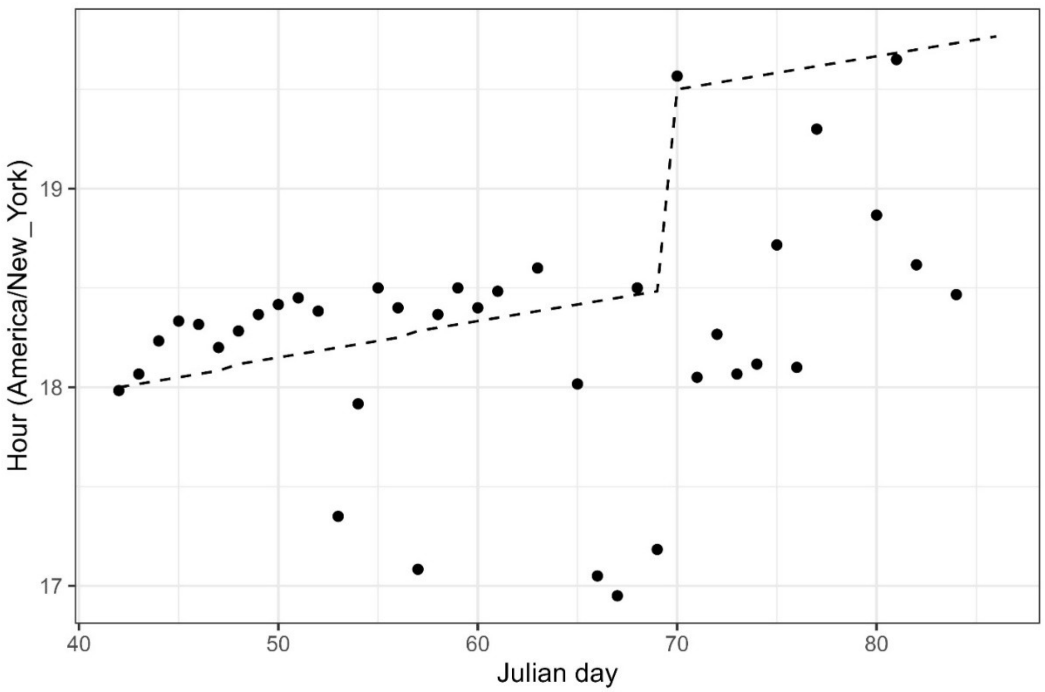


Figure S6. Per-night estimates of roost onset in relation to sunset (dashed line) by doy for the American tree sparrow.

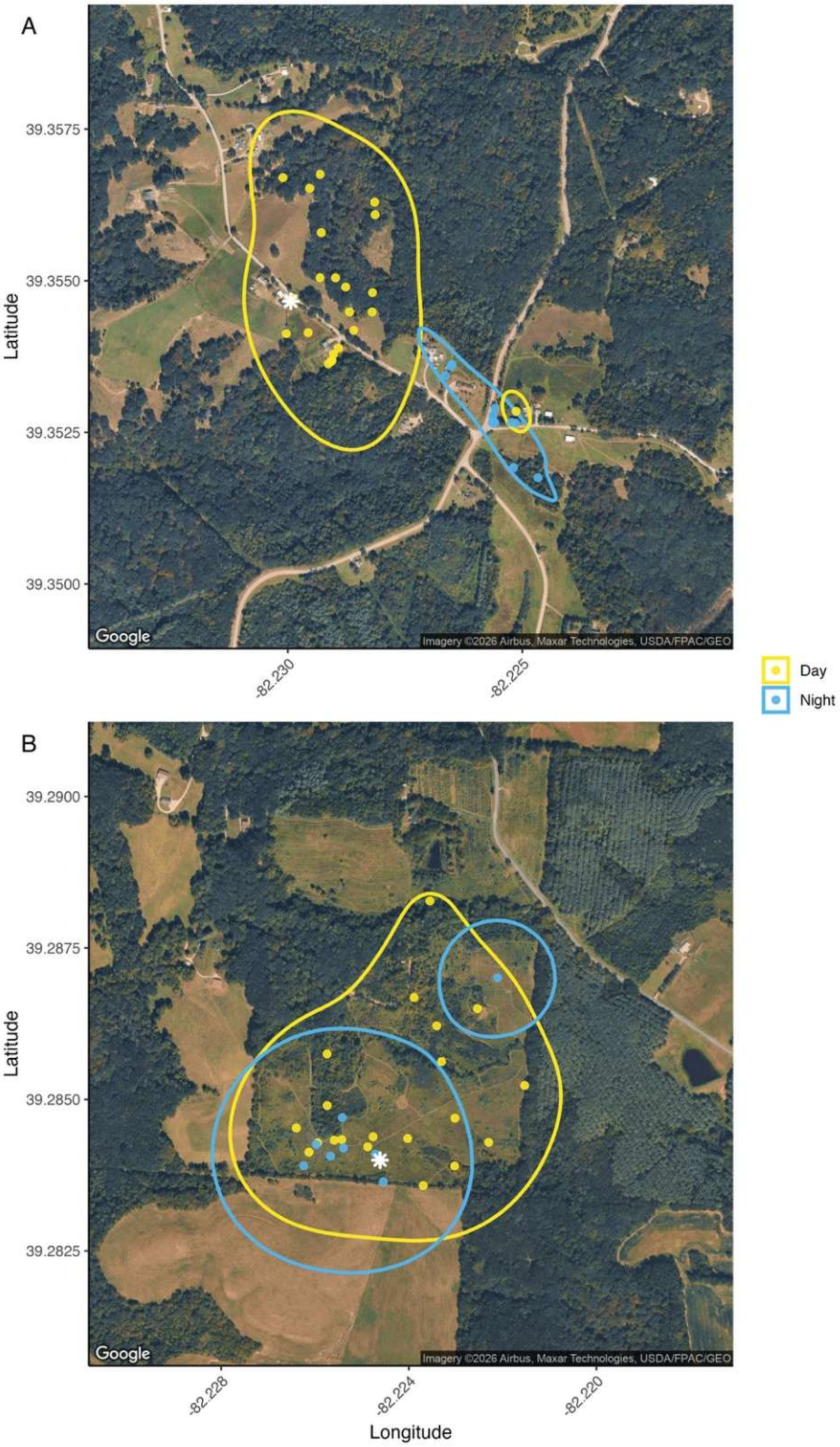


Figure S7: Diurnal (yellow) and roost site (blue) kernel density ranges (polygons) and manual telemetry detections (dots) for A) Dark-eyed Junco 52550 and B) American Tree Sparrow 52491. White stars indicate the location of the Motus receiving station at the field site. Manual telemetry was conducted as part of a larger study (see Morgan and Williams 2025).
